# MaskTalk: cell-identity-gated spatial lag for target-aware cell-cell communication inference in high-resolution spatial transcriptomics

**DOI:** 10.64898/2026.08.02.741948

**Authors:** Panpan Jia, Liqiang Chu, Zhenhua Ren, Hongwei Cui, Bin Shao

## Abstract

**Motivation:** High-resolution spatial transcriptomics enables single-cell ligand-receptor analysis, but unmasked receptor spatial lags include receptor expression from non-target neighbors, complicating the attribution of local communication signals to specified source-target cell-type pairs.

**Results:** We present MaskTalk, a Python package implementing the cell-identity-gated spatial lag model (CIG-SLM). CIG-SLM restricts receptor-side neighborhoods to target cells through **W**^(*C*)^ = **WD**^(*C*)^. In cell-level breast cancer Visium HD data, CIG-SLM produced target-cell-dependent communication profiles relative to the matched LIANA+ bivariate unmasked baseline, and masked-specific records showed larger between-condition effect sizes and shorter physical source-target distances. Public breast cancer Xenium data demonstrated that MaskTalk runs on external cell-level spatial data and exhibits target-aware masking behavior.

**Availability and Implementation:** Implemented in Python with AnnData; code is available at https://github.com/JiaPP1994/MaskTalk. Software and data archive DOIs are 10.5281/zenodo.21735883 and 10.5281/zenodo.21735951, respectively.

**Supplementary Information:** Supplementary Methods S1-S4, Figures S1-S3, and Tables S1-S5.

## 1 Introduction

High-resolution spatial transcriptomics technologies are advancing tissue microenvironment analysis from low-resolution spots to single-cell or near-single-cell resolution. New-generation technologies, such as Visium HD, Xenium, and other imaging-based or sequencing-based spatial transcriptomics platforms, retain gene expression information while recording the in situ coordinates and physical proximity of cells (Janesick *et al*. 2023, Oliveira *et al*. 2025). In tissues with pronounced spatial heterogeneity, such as breast cancer tissues, previous single-cell and spatially resolved studies have shown that tumor cell states, stromal organization, and the immune microenvironment are highly intertwined (Wu *et al*. 2021, Wang *et al*. 2024). After cell segmentation, these data provide a new basis for local ligand-receptor communication inference at single-cell resolution.

Existing cell-cell communication inference methods can be broadly grouped by their core statistical unit into methods based on cell-population expression aggregation and bivariate statistical methods that explicitly use spatial coordinates. Classical implementations of the former were developed primarily for scRNA-seq data. CellChat and CellPhoneDB typically aggregate ligand and receptor expression using predefined cell populations and output potential communication probabilities, interaction means, or statistical significance at the level of cell-population pairs (Efremova *et al*. 2020, Jin S *et al*. 2021, Jin S *et al*. 2025, Troule *et al*. 2025). NicheNet further links candidate ligands to downstream target gene responses in receiver cells (Browaeys *et al*. 2020). In these methods, the source and target are generally specified by predefined cell-population labels. Spatial bivariate methods such as SpatialDM and LIANA+ use spatial coordinates to detect coordinated variation between ligand expression and neighborhood receptor expression; local Moran’s *R* links ligand expression at a source location to the receptor spatial lag (Li Z *et al*. 2023, Dimitrov *et al*. 2024). A key question in spatial bivariate communication inference is therefore which neighborhood receptor signals should enter the local statistic for the current source cell.

In unmasked implementations of spatial bivariate statistics, receptor expression from all neighboring cells contributes to the receptor spatial lag, without screening receptor neighbors by a specified target cell type (Li Z *et al*. 2023, Dimitrov *et al*. 2024). In tissues with highly heterogeneous cell types, the neighborhood of a source cell may contain both the prespecified target cells and multiple non-target cell populations. Receptor expression in non-target cells can form a receiver-side bystander signal, making it difficult to exclude the influence of non-target neighbors when local Moran’s *R* is summarized for a source-target cell-type pair. Once non-target receptor expression enters the receptor spatial lag, subsequent grouping can only change output labels and cannot redefine the receptor sources actually used by the local statistic.

This issue is less prominent in low-resolution spot data because one spot often contains multiple cells, and cell identity usually enters the analysis as cell-type proportions, deconvolution results, spot groups, or spatial-domain labels. With the emergence of cell-level spatial data from platforms such as Visium HD and Xenium, receptor-side neighbors can be explicitly screened at single-cell resolution. However, existing spatial bivariate statistics generally use cell type for result stratification or candidate-pair definition rather than reconstructing the receptor spatial lag actually used by the local statistic.

Existing spatial omics methods use spatial information and cell identity at different stages of modeling. Squidpy provides infrastructure for spatial adjacency graphs and spatial autocorrelation analysis (Palla *et al*. 2022). COMMOT, SpaTalk, Giotto, stLearn, NICHES, MISTy, Niche-DE, NCEM, MESSI, and SEGCECO characterize spatial communication and microenvironmental effects from the perspectives of optimal transport, spatial neighborhoods, cell-type pairs, signal proxies, multiview neighborhood modeling, niche-associated expression, spatial graph modeling, or graph representation learning (Dries *et al*. 2021, Li D *et al*. 2021, Shao *et al*. 2022, Tanevski *et al*. 2022, Cang *et al*. 2023, Fischer *et al*. 2023, Pham *et al*. 2023, Raredon *et al*. 2023, Mason *et al*. 2024, Vasighizaker *et al*. 2024). These methods provide the relevant methodological context for spatial communication analysis. MaskTalk further focuses on the key problem of receiver attribution in cell-level local ligand-receptor statistics. Before calculating the statistic, the specified target-cell identity is written directly into the receptor spatial lag operator so that the receptor-side signal source used by the local statistic is consistent with the target cell type specified by the research question. The conceptual positioning of related methods is provided in Supplementary Table S5.

Here, we propose the cell-identity-gated spatial lag model (CIG-SLM) and implement it in the Python package MaskTalk. For a given target cell type, CIG-SLM constructs an identity diagonal matrix and right-multiplies the spatial adjacency matrix by it, so that the receptor spatial lag receives contributions only from cells that are both physically adjacent and of the specified target cell type. The main contributions of MaskTalk are as follows: it incorporates target-cell identity into the receiver-side spatial lag operator before local Moran’s *R* calculation; it provides an end-to-end workflow for spatial graph construction, receiver-side gated ligand-receptor inference, a matched unmasked baseline, local permutation tests, aggregation at the sample, subject, and condition levels, and visualization; and it uses cell-level breast cancer Visium HD data and public Xenium data to demonstrate how target-aware masking affects local statistics, source-target attribution, and output networks.

## 2 Materials and methods

### 2.1 Input data, cell-type annotation, and spatial graph construction

MaskTalk uses an AnnData object as input. The expression matrix is obtained from adata.X or adata.raw.X, cell-type annotations from adata.obs, and spatial coordinates from adata.obsm[“spatial”]. Multisample data must contain Sample_ID, Subject_ID, and Condition_ID. Spatial graphs are constructed separately by Sample_ID to avoid spurious adjacency edges between different tissue sections.

MaskTalk supports spatial contact and kNN graphs. A spatial contact graph is based on Delaunay triangulation and can be pruned using a distance threshold; kNN graphs construct spatial neighborhoods at different scales by specifying the number of nearest neighbors. The validation analyses used both spatial contact and spatial kNN graphs. Detailed parameters and workflows are provided in the Supplementary Methods.

### 2.2 Ligand-receptor resources and local bivariate spatial statistics

This study used the consensus ligand-receptor resource and local bivariate Moran’s *R* for local communication statistics. In mt.tl.communication(), resource_name and local_name specify the ligand-receptor resource and local statistic, respectively, with defaults of consensus and morans. The examples, Visium HD validation, and public Xenium validation all used these settings.

CIG-SLM reconstructs the spatial weight matrix before calculating the local bivariate statistic, so that target-cell identity first enters the receptor spatial lag. Complex-expression aggregation, basic expression filtering, local Moran’s *R* , and local permutation calculations reuse the LIANA+ bivariate interface (Dimitrov *et al*. 2024). The masked and matched unmasked modes use the same computational rules; the gated matrix **WD**^(*C*)^ and the unmasked matrix **W** are each scaled according to the SpatialDM weight-normalization rule used by this interface before entering the statistic calculation (Li Z *et al*. 2023).

### 2.3 Cell-identity-gated spatial lag model

For a spatial transcriptomics sample containing *n* cells, define the spatial adjacency matrix **W** *∈* ℝ^*n*×*n*^. Here, *W*_*ij*_ denotes the spatial graph weight directed from source cell *i* to receiver-side cell *j*; in a binary adjacency graph, *W*_*ij*_ = 1 indicates that the two cells are adjacent, whereas *W*_*ij*_ = 0 indicates that they are not adjacent.

For target cell type *C*, construct the binary identity vector:

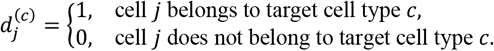

This vector is written into the identity diagonal matrix:

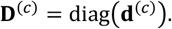

The target-masked spatial adjacency matrix is defined as:

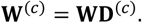

Matrix multiplication gives:

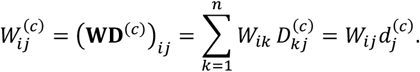

Because 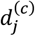 depends only on the receiver-side index *j*, right multiplication by **D**^(*C*)^ is equivalent to multiplying the entire *j*, th column of **W** by 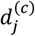. This operation retains only the original adjacency weights corresponding to the specified target receptor cells, thereby implementing receiver-side column gating and allowing target-cell identity to directly restrict the signal source of the receptor spatial lag.

### 2.4 Receptor spatial lag, local permutation test, and summary metrics

The unmasked receptor spatial lag receives contributions from all neighboring cells:

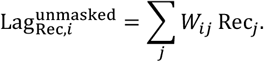

After target masking:

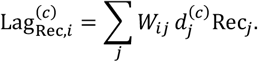

Here, Rec_*j*_ denotes receptor expression in receiver-side cell *j*. For source cell *i*, MaskTalk uses ligand expression Lig_*i*_ and the target-masked receptor spatial lag Lag^(*C*)^ as the two inputs to local bivariate Moran’s *R*_*i*_ and obtains a local *P* value through a local permutation test. Within-sample summary metrics include Mean_Morans_R, Local_sig_cell_count, Total_cell_count, and Local_sig_cell_prop; the latter is abbreviated as Sig prop.

### 2.5 Matched LIANA+ bivariate unmasked baseline and ablation comparison design

The matched LIANA+ bivariate unmasked baseline uses the same expression input, spatial graph, ligand-receptor resource, local statistic, and permutation parameters as the masked mode but does not use *D*^(*C*)^. Candidate target cell types are used only for result matching and stratification and do not enter the spatial adjacency matrix or receptor spatial lag calculation. This baseline constitutes a direct ablation control for receiver-side identity gating in CIG-SLM.

### 2.6 Visium HD validation data and public Xenium data

The Visium HD validation used a de-identified cell-level AnnData object from breast cancer that contained the expression matrix, spatial coordinates, cell-type annotations, and grouping information required for MaskTalk validation. Breast cancer was selected as the validation setting because previous studies have shown pronounced heterogeneity in cell states and the spatial microenvironment, making it suitable for evaluating high-resolution spatial communication methods (Wu *et al*. 2021, Wang *et al*. 2024).

The public validation used In Situ Sample 1, Replicate 1 from the 10x Genomics Human Breast Xenium preview dataset and the supervised annotation provided on the 10x data page (Janesick *et al*. 2023). This module served as an external public-data validation to assess the software’s ability to run on external cell-level spatial data and its target-aware masking behavior.

### 2.7 Condition comparison, mode-specific records, and distance statistics

For a grouped multisample design, mt.tl.aggregate_statistics() aggregates within-sample communication metrics by Subject_ID, Condition_ID, spatial graph, source cell type, target cell type, and ligand-receptor pair, using Subject_ID as the independent replicate. It then calculates group means, Mann-Whitney U test *P* values, and Benjamini-Hochberg FDR between condition1 and condition2. The direction of the difference is fixed as condition2 - condition1.

Communication records that pass the prespecified effect-size thresholds are termed threshold-positive LR edges. The mode-specific analysis focuses on masked-specific and unmasked-specific records and compares their target-cell lineage distributions, |*Δ*Moran′s *R*| , and physical source-target distances. Physical source-target distance is defined as the Euclidean distance between the spatial centroids of the source and target cell types, calculated from the adata.obsm[“spatial_um”] coordinates. Representative ligand-receptor axes were selected jointly on the basis of target-dependent profiles, effect-size rankings, graphical readability, and known context to show typical algorithmic behavior.

### 2.8 Software implementation and reproducibility

MaskTalk comprises the pp, tl,and pl modules, which are responsible for spatial graph construction, communication inference and statistical aggregation, and network and bubble-plot visualization, respectively. The main APIs include mt.pp.build_spatial_graph(), mt.tl.communication(), mt.tl.aggregate_statistics(), mt.pl.network(), and mt.pl.bubble(). The examples, benchmarks/demo_validation, and benchmarks/xenium_janesick directories correspond to the basic examples, Visium HD algorithm validation, and public Xenium validation, respectively.

## 3 Results

### 3.1 CIG-SLM incorporates target-cell identity into the receiver-side spatial lag

MaskTalk jointly inputs the expression matrix, cell-type annotations, and spatial graph into CIG-SLM (Figure 1A). A conventional unmasked model includes receptor expression from all neighbors around a source cell in the receptor spatial lag; when the research question concerns a specific target cell type, non-target neighbors form a receiver-side bystander signal (Figure 1B). CIG-SLM constructs an identity diagonal matrix for each candidate target cell type and right-multiplies the spatial weight matrix by it, **W**^(*C*)^ = **WD**^(*C*)^. This receiver-side column gating filters the columns of the spatial weight matrix by target-cell identity and retains only the original adjacency weights corresponding to the specified target receptor cells. In other words, CIG-SLM changes the object to which the receptor signal is attributed at the spatial-lag level (Figure 1C).

**Figure 1.**
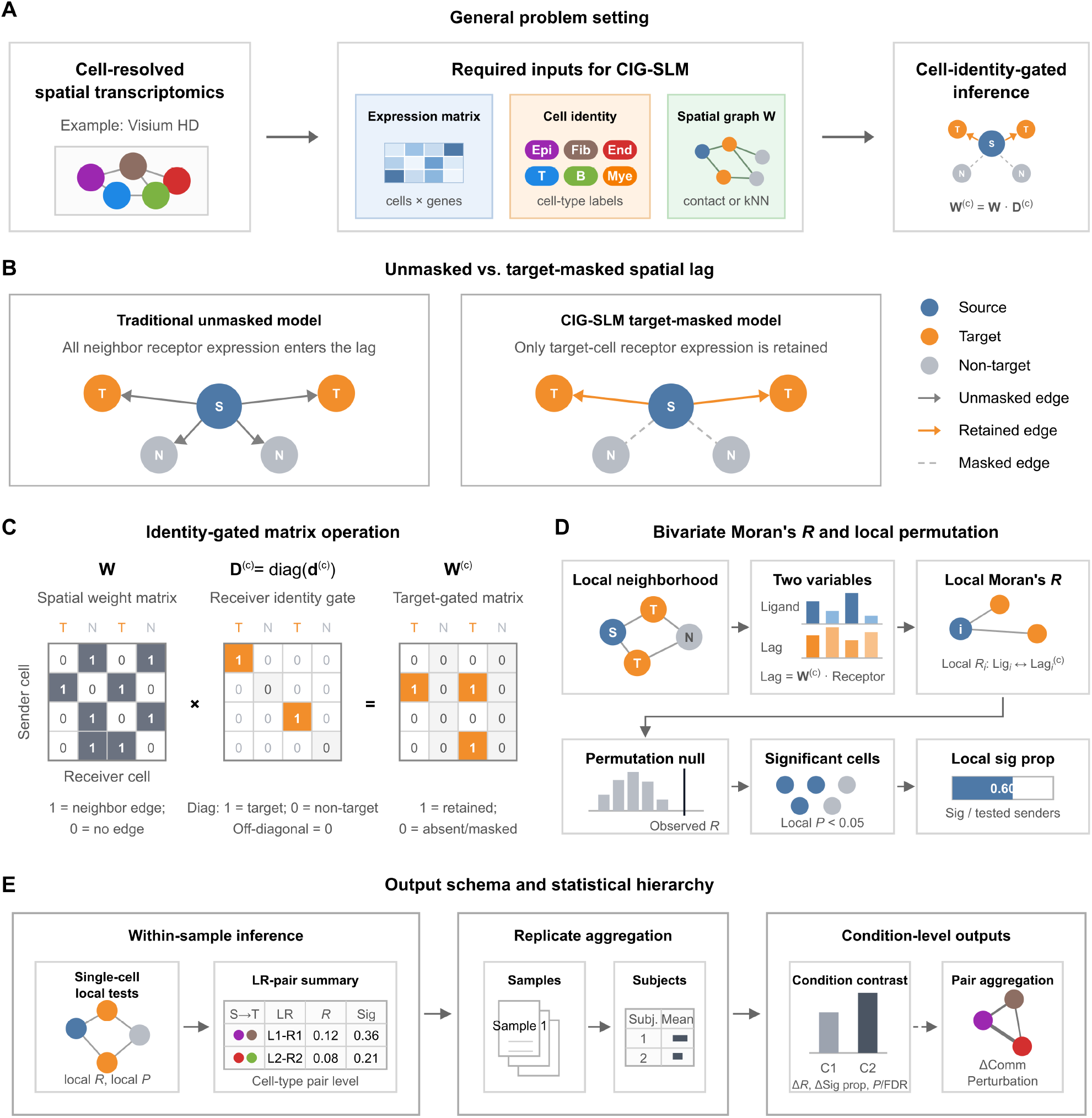
CIG-SLM principles and statistical output levels in MaskTalk. **(A)** CIG-SLM integrates the expression matrix, cell-type annotations, and spatial graph W, and constructs **W**^(*C*)^ = **WD**^(*C*)^ through **D**^(*C*)^. **(B)** The unmasked spatial lag summarizes receptor expression from all neighbors, whereas CIG-SLM retains signals only from neighbors of the specified target cell type. **(C)** Right multiplication by the diagonal matrix implements receiver-side column gating and filters receptor-side adjacency-matrix columns by the specified target-cell identity. **(D)** Ligand expression and 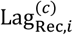 are input into local bivariate Moran’s *R*, and a local permutation test yields the local *P* value and Sig prop. **(E)** MaskTalk outputs single-cell local statistics, within-sample summaries, and optional comparisons at the subject and condition levels. *Alt text: Multipanel schematic showing how MaskTalk gates the spatial weight matrix by target-cell identity and calculates local Moran’s R and multilevel communication results from the gated receptor spatial lag*.

### 3.2 MaskTalk outputs cell-level local communication tistics and multilevel summaries

On the gated spatial graph, MaskTalk inputs source-cell ligand expression and 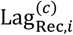 into local bivariate Moran’s *R* and obtains a local *P* value through a local permutation test (Figure 1D). The software then summarizes Mean_Morans_R, Local_sig_cell_count, Total_cell_count, and Sig prop. For grouped multisample designs, MaskTalk further aggregates results at the sample, subject, and condition levels to generate condition-comparison and cell-pair results (Figure 1E).

### 3.3 Target masking reveals target-cell-dependent communication profiles

We used the unmasked LIANA+ bivariate analysis as a matched baseline representing the existing local bivariate implementation and compared CIG-SLM with the matched LIANA+ bivariate unmasked baseline in the Visium HD validation data. For the same source type, ligand-receptor pair, and spatial graph, the baseline retains the unmasked receptor spatial lag of LIANA+ bivariate, with candidate target cell types used only for result matching. CIG-SLM instead incorporates target-cell identity into the receptor spatial lag through **D**^(*C*)^, thereby producing a target-cell-dependent profile.

For example, with Endo-4 as the source type, SPARC→ENG as the ligand-receptor axis, and a spatial kNN (*k*=15) graph, the unmasked curve was approximately flat across candidate target cell types, whereas the masked curve peaked at candidate target cells including Endo-1 and Endo-4 (Figure 2A). Supplementary Figure S1A-C further shows that this pattern was reproduced for SPARC→ENG in Endo-3 with a spatial contact graph, SPP1→CD44 in Fibro-5 with a spatial contact graph, and APOE→LRP1 in Mye-5 with a spatial kNN (*k*=15) graph. This result is consistent with the expectation of CIG-SLM: the same source cell type and ligand-receptor pair produce different target profiles only when target-cell identity enters the receiver-side spatial lag.

**Figure 2.**
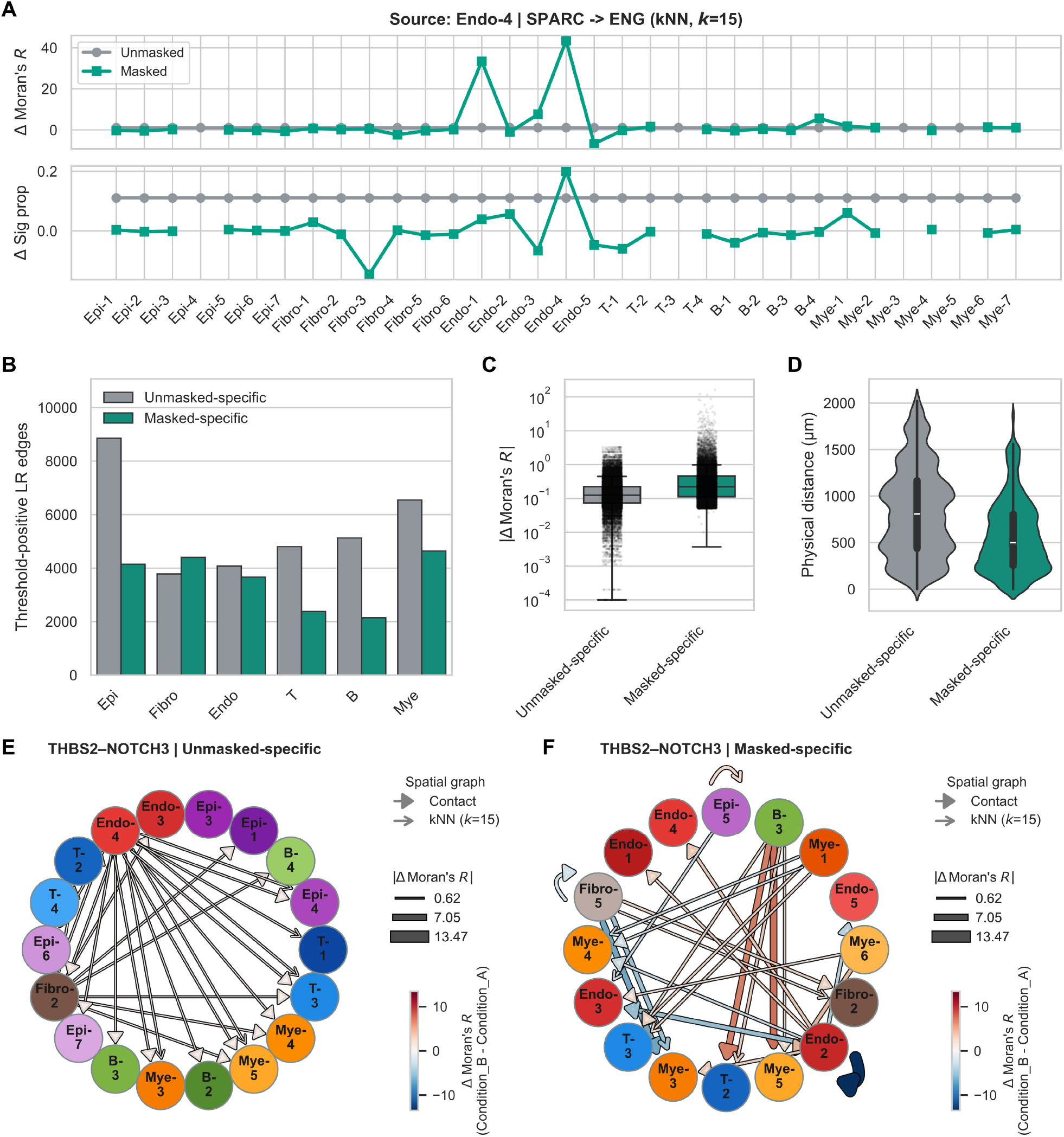
Target masking reveals target-cell-dependent communication profiles and mode-specific signals. **(A)** Bystander validation for SPARC→ENG with Endo-4 as the source type on a spatial kNN (*k*=15) graph. Gray indicates the matched unmasked baseline and green indicates the masked mode. **(B)** Numbers of threshold-positive LR edges among different target-cell lineages for unmasked-specific and masked-specific records. **(C)** Distributions of |*Δ*Moran′s *R*| for the two record classes. **(D)** Distributions of physical distances between the spatial centroids of source and target cell types. **(E**,**F)** Unmasked-specific and masked-specific communication networks for THBS2→NOTCH3; nodes denote cell types, edge width denotes |*Δ*Moran′s *R*|, edge color denotes *Δ*Moran′s *R*, and arrow style denotes spatial graph type. *Alt text: Multipanel software workflow showing the end-to-end MaskTalk process from AnnData input through spatial graphs and CIG-SLM inference to summary tables, network plots, and bubble plots*.

### 3.4 Masked-specific and unmasked-specific records show different signal characteristics

We defined records that reached the prespecified effect-size threshold only in the masked mode as masked-specific and those that reached the threshold only in the matched unmasked baseline as unmasked-specific. We then compared the lineage distributions, between-condition effect sizes, and physical source-target distances of the two record classes. The Visium HD validation data yielded 21,374 masked-specific and 33,194 unmasked-specific records. Masked-specific records had a higher median |*Δ*Moran′s *R*| (0.2206 vs. 0.1245) and a shorter median physical distance (499.35 μm vs. 808.23 μm; record-level descriptive Mann-Whitney U test, *P*<0.001; Figure 2B-D). These results indicate that receiver-side identity gating changes the effect-size and spatial-distance structures of records that pass the threshold, rather than merely reducing the number of records.

To examine whether this trend depended on the mode-specific definition, we performed a sensitivity analysis using all records that passed the threshold. All masked records likewise showed a higher median |*Δ*Moran′s *R*| (0.2894 vs. 0.1237) and a shorter median physical distance (492.62 μm vs. 570.47 μm; Supplementary Figure S2A-C). This analysis compares the outputs of the two computational modes at the record level.

### 3.5 Representative ligand-receptor axes show different source-target connection structures

At the ligand-receptor-axis level, mode-specific records also showed different source-target connection structures. For THBS2→NOTCH3, the unmasked-specific network mainly showed positive edges diverging from a small number of source-cell states to multiple target-cell states. The masked-specific network retained more specific source-target pairs among cell states labeled Endo-, B-, Mye-, and Fibro-, and included effects with different directions and larger magnitudes (Figure 2E,F). Similar changes were observed for SEMA4C→PLXNB2 and COL1A1→CD44: unmasked-specific edges often connected a small number of source states broadly to multiple target states, whereas masked-specific edges were redistributed to more specific source-target pairs and showed more heterogeneous effect directions and magnitudes (Supplementary Figure S2D-G).

### 3.6 Public Xenium data validate target-aware masking behavior

To validate software compatibility and target-aware masking behavior on public data, we used the public Xenium data described above. This object contained 167,780 cells, 313 genes, and 20 supervised cell types.

For the same source type, ligand-receptor pair, and spatial graph type, the target-profile ranges of both Mean Moran’s *R* and Sig prop were 0 for the matched unmasked baseline, whereas the masked mode showed variation dependent on the candidate target cell type (Supplementary Figure S3A,B). For MMRN2→CLEC14A in Endothelial cells on a spatial contact graph, the target-profile ranges of masked Mean Moran’s *R* and Sig prop were 107.06 and 0.770, respectively, whereas both unmasked ranges were 0 (Supplementary Figure S3D). This result shows that the target-cell gating behavior of CIG-SLM can be reproduced on an external public-data object.

### 3.7 MaskTalk provides reproducible software workflows and examples

MaskTalk packages the model and validation analyses into end-to-end Python workflows. Starting from an AnnData object, users can sequentially call mt.pp.build_spatial_graph(), mt.tl.communication(), and mt.tl.aggregate_statistics() to construct spatial graphs, perform communication inference, and summarize results at the sample, subject, and condition levels (Figure 3A). mt.pl.network() and mt.pl.bubble() generate network and bubble plots (Figure 3B-D).

**Figure 3.**
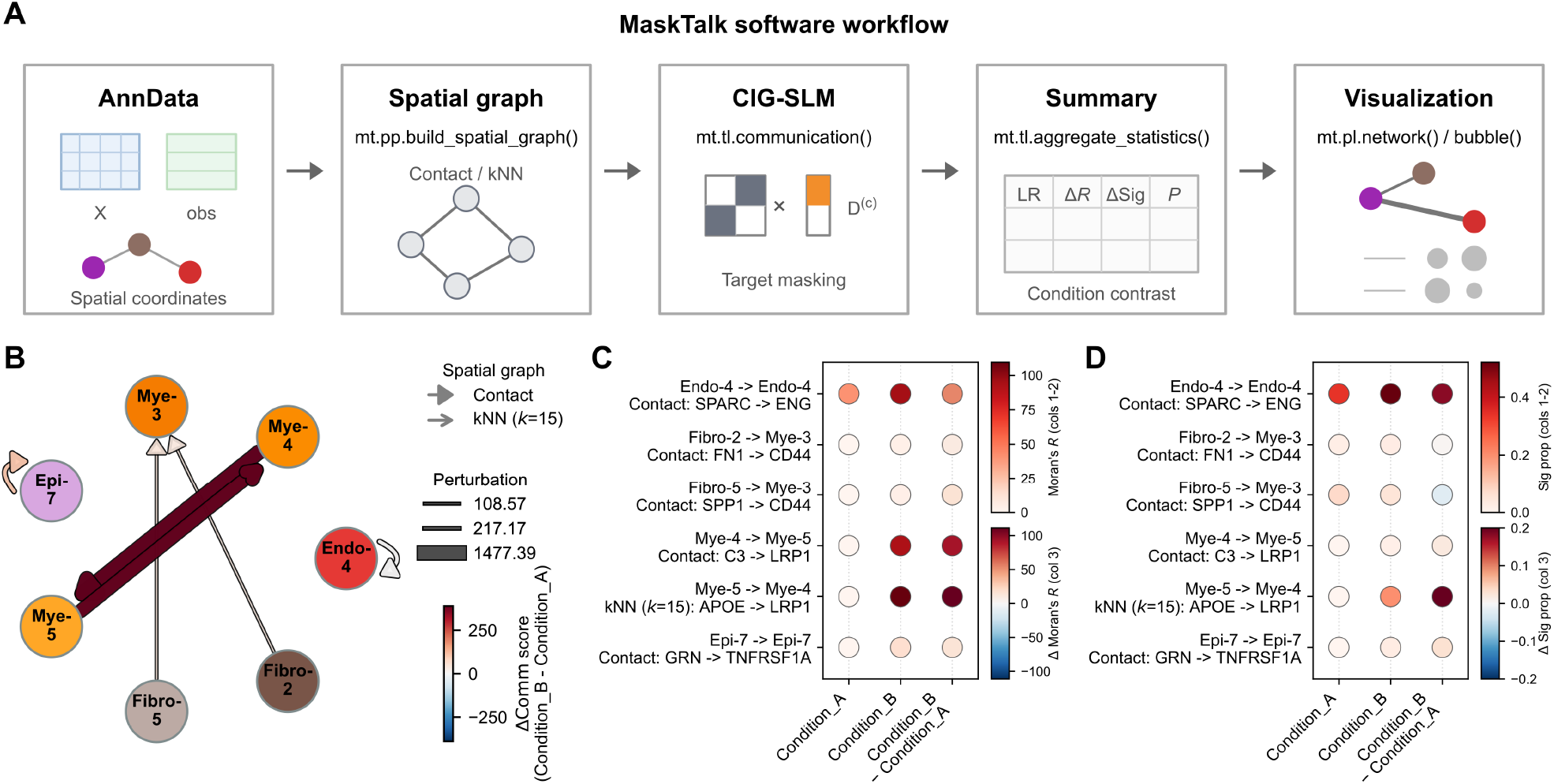
End-to-end target-masked spatial communication workflow in MaskTalk. **(A)** From AnnData input through spatial graph construction, CIG-SLM inference, statistical summarization, and visualization. **(B)** Cell-pair communication network in the example data; nodes denote cell types, edge width denotes Perturbation, edge color denotes the between-condition *Δ*Comm score, and arrow style denotes spatial graph type. **(C**,**D)** Ligand-receptor bubble plots showing between-condition changes in selected cell pairs and ligand-receptor pairs using Moran’s *R* and Sig prop. *Alt text: Multipanel results comparing the matched unmasked baseline and masked mode, showing target-cell-dependent curves, lineage counts, effect sizes, and distance distributions for mode-specific records, and the source-target connection structure of THBS2→NOTCH3*.

The examples, benchmarks/demo_validation, and benchmarks/xenium_janesick directories in the repository correspond to basic usage, Visium HD algorithm validation, and public Xenium validation, respectively. Figure-source data, processed results, and checksum files are provided with the data archive to enable inspection of algorithm outputs.

## 4 Discussion

The core contribution of MaskTalk is to incorporate target-cell identity as a matrix-gating term in the receiver-side spatial lag, allowing target-cell identity to participate directly in the calculation of local spatial statistics. This design changes the signal sources of the receptor spatial lag so that the receptor-side input to local Moran’s *R* is consistent with the specified target cell type. The Visium HD validation data showed that CIG-SLM can produce target-dependent communication profiles and alter the effect sizes, distance distributions, and source-target connection structures of records that pass the threshold. Public Xenium validation showed that the implementation can run on external cell-level spatial data and produce the expected target-aware masking behavior.

Existing spatial cell-cell communication methods have different analytical objectives and output definitions. CIG-SLM modifies the receiver-side spatial lag before local Moran’s *R* calculation; therefore, the existing comparator that most directly corresponds to its target quantity is the unmasked implementation of LIANA+ bivariate. We accordingly constructed a matched LIANA+ bivariate unmasked baseline and compared the effects of receiver-side target-cell identity gating on target profiles and output structures while holding the expression input, ligand-receptor resource, spatial graph, local statistic, and permutation parameters constant. Conceptual and functional comparisons with other spatial communication methods are provided in Supplementary Table S5.

MaskTalk is an end-to-end software workflow for target-aware local bivariate ligand-receptor inference and can independently perform analyses from spatial graph construction and receiver-side gated inference to multilevel summarization and visualization. It reuses established ligand-receptor resources and bivariate spatial-statistical interfaces, while the added CIG-SLM advances cell identity from a result-grouping variable to a component of the spatial lag operator before local-statistic calculation. Unlike MISTy, Niche-DE, and NCEM, which focus on spatial context, niche composition, or neighborhood predictive relationships, MaskTalk addresses receiver-side target-cell attribution in local ligand-receptor statistics for high-resolution, cell-level tissues with highly heterogeneous cell types (Tanevski *et al*. 2022, Fischer *et al*. 2023, Mason *et al*. 2024).

Interpretation of MaskTalk results should consider cell-segmentation quality, cell-type annotation accuracy, spatial graph construction, ligand-receptor resources, and gene-detection sensitivity. Errors in cell segmentation and transcript assignment affect the cell-level expression matrix, spatial neighborhood definition, and downstream analysis (Jin K *et al*. 2025). The method identifies candidate ligand-receptor associations jointly supported by ligand expression, receptor expression, and spatial proximity, providing candidates for subsequent validation of protein binding and functional signaling.

Future extensions of MaskTalk may include richer spatial graphs and validation modules. Histology-aware graphs could use tissue images or morphological features to adjust adjacency edge weights; cell-boundary contact graphs could replace centroid proximity with actual boundary contact when segmentation boundaries are available; ligand-specific distance kernels could assign different distance-decay weights to different ligands; and simulated ground-truth benchmarks and additional public datasets could be used to evaluate the sensitivity and robustness of the gating strategy. In data with spatial protein or cell-boundary information, receptor protein abundance and membrane-contact constraints could also be incorporated to support interpretation of candidate communication axes.

## Author contributions

Panpan Jia: Conceptualization, Methodology, Software, Formal analysis, Data curation, Validation, Visualization, Writing – original draft, Writing – review & editing, and Project administration. Liqiang Chu: Formal analysis, Investigation, Validation, and Writing – review & editing. Zhenhua Ren: Resources, Validation, Supervision, and Writing – review & editing. Hongwei Cui: Funding acquisition, Resources, Validation, and Writing – review & editing. Bin Shao: Supervision, Funding acquisition, Project administration, and Writing – review & editing. All authors reviewed and approved the final manuscript.

## Funding

This work was supported by the National Natural Science Foundation of China [grant number 12374406] and the Oncology Discipline Development Program of Inner Mongolia Medical University [grant number YKD2024XK004].

## Conflict of interest

None declared.

## Data availability

Example data, validation data, processed results, and figure-source data are available through the Zenodo data record, with version DOI 10.5281/zenodo.21735951 and concept DOI 10.5281/zenodo.20812932. The raw public Xenium data are available from the 10x Genomics preview dataset page and correspond to the study by Janesick et al. (Janesick *et al*. 2023). The specific download page is provided in Supplementary Methods S1.2, and the Zenodo archive provides processed Xenium validation outputs and checksum files.

## Code availability

The MaskTalk source code, installation instructions, API documentation, example workflows, benchmark workflows, and test code are available at https://github.com/JiaPP1994/MaskTalk. The Software Zenodo version DOI is 10.5281/zenodo.21735883, and the concept DOI is 10.5281/zenodo.20814048.

## Ethics statement

The Visium HD validation data were obtained from a study approved by the Ethics Committee of Peking University Cancer Hospital (approval number 2025KT121), and written informed consent was obtained from all participants in the original study. This study used only de-identified data for software validation and involved no new participant recruitment or intervention. The public Xenium validation used publicly available de-identified data.

## Acknowledgements

During manuscript preparation, the authors used ChatGPT and Kimi to assist with Chinese-English translation, language editing, and code review. All related outputs were reviewed by the authors. The final manuscript and software implementation were approved by the authors, who take responsibility for both.

## Notes

https://github.com/JiaPP1994/MaskTalk

https://doi.org/10.5281/zenodo.21735883

https://doi.org/10.5281/zenodo.21735951

### Competing Interest Statement

The authors have declared no competing interest.

