## Supplementary Materials for "MaskTalk: cell-identity-gated spatial lag for target-aware cell-cell communication inference in high-resolution spatial transcriptomics"

### Contents

Supplementary Methods: S1-S4

Supplementary Figures: Figures S1-S3

Supplementary Tables: Tables S1-S5

### Supplementary Methods

#### S1 Data sources and preprocessing

##### S1.1 Visium HD validation data

The Visium HD validation used a de-identified cell-level AnnData object constructed from a breast cancer Visium HD spatial transcriptomics study. The original study was approved by the Ethics Committee of Peking University Cancer Hospital (approval number 2025KT121), and written informed consent was obtained from all participants. The upstream cell-level object was constructed by nucleus-based expansion: cell nuclei were first identified using StatDist, and nearby transcripts were then assigned by outward spatial expansion. Subsequent MaskTalk analyses used cells as the basic spatial units.

The object retained the expression matrix, spatial coordinates, cell-type annotations, Sample_ID, Subject_ID, and Condition_ID required by MaskTalk, while sensitive clinical identifiers and identifiable sample information were removed. MaskTalk can use a spatial graph already present in AnnData or construct a spatial contact or kNN graph from spatial coordinates through mt.pp.build_spatial_graph(). The examples workflow in this study constructs spatial graphs from this object, whereas the demo_validation workflow uses a derived object containing spatial graphs. The object and its derived results were used for the basic examples workflow and for matched unmasked and masked ablation, bystander validation, mode-specific signal characterization, and all-threshold-positive sensitivity analysis in the demo_validation benchmark. The example object, processed outputs, and figure-source data are available through the Zenodo data record.

##### S1.2 Public Xenium data

The public validation module used In Situ Sample 1, Replicate 1 from the 10x Genomics Human Breast Xenium preview dataset. The raw data and Cell_Barcode_Type_Matrices.xlsx were downloaded from the 10x Genomics preview dataset page: https://www.10xgenomics.com/products/xenium-in-situ/preview-dataset-human-breast. For the analysis, cell_groups.csv was exported from the Xenium R1 Fig1-5 (supervised) sheet of Cell_Barcode_Type_Matrices.xlsx and integrated with the Xenium output files into an AnnData object.

The external raw files used by the workflow included Cell_Barcode_Type_Matrices.xlsx, cell_feature_matrix.h5, cells.parquet, and gene_panel.json. The prepared Xenium object contained 167,780 cells, 313 genes, and 20 supervised cell types. This module was a single-sample external validation that demonstrated the ability of MaskTalk to run on public Xenium data and the algorithmic behavior of target-aware masking across candidate target cell types.

##### S1.3 AnnData fields and metadata

MaskTalk uses an AnnData object as input. The recommended input contains an expression matrix, cell-type annotations, spatial coordinates, and optional grouping metadata. When an externally precomputed spatial graph is used, the spatial adjacency matrix should be stored as a sparse matrix in AnnData and passed to the communication inference function through Graph_key or the corresponding parameter. For multisample data, spatial graphs are constructed or read separately by Sample_ID to avoid spurious adjacency between different tissue sections. In clinical or experimental replicate designs, Subject_ID denotes the independent statistical replicate. If each sample is itself an independent replicate, Subject_ID and Sample_ID may refer to the same field.

MaskTalk input fields, output fields, and definitions of the main statistical metrics are provided in Table S1.

### S2 Spatial graphs and communication inference

#### S2.1 Spatial contact and kNN graph construction

MaskTalk supports spatial contact graphs, kNN graphs, and user-provided external spatial graphs. A spatial contact graph is based on Delaunay triangulation, with optional distance-threshold pruning of excessively long edges; a kNN graph is constructed using a specified number of nearest neighbors. The examples and validation analyses in this study used $k$=15 for kNN graphs. Spatial graphs are stored as sparse matrices in AnnData and used as spatial weight matrices for subsequent receptor spatial lag calculation.

#### S2.2 Ligand-receptor resources and local bivariate statistical parameters

MaskTalk places CIG-SLM before local bivariate statistic calculation. By reconstructing the receiver-side spatial weight matrix, target-cell identity directly restricts the signal source of the receptor spatial lag. The examples and validation workflows in this study used the consensus ligand-receptor resource and Moran's $R$ as the local spatial statistic. The expression matrix used for inference should contain normalized expression, such as log-normalized expression, rather than scaled z-scores centered by gene and divided by the standard deviation. Complex-expression representation, ligand-receptor resource parsing, basic expression filtering, local Moran's $R$, and local permutation calculations reuse the LIANA+ bivariate interface.

MaskTalk passes either the unmasked matrix $\mathbf{W}$ or the gated matrix $\mathbf{W}^{(c)} = \mathbf{W}\mathbf{D}^{(c)}$ to the same LIANA+ bivariate computational workflow. For either input spatial weight matrix $\mathbf{A} \in \{\mathbf{W}, \mathbf{W}^{(c)}\}$, the interface applies global normalization according to the SpatialDM rule:

$$\widetilde{\mathbf{A}} = \frac{n\mathbf{A}}{\sum_{i,j} A_{ij}},$$

where $n$ is the number of cells and the tilde denotes the normalized spatial weight matrix; this scaling sets the sum of the matrix weights to $n$. The masked and matched unmasked modes use the same local-statistic implementation and normalization rule. Their prespecified difference at the input level is the CIG-SLM reconstruction of the receiver-side weight matrix.

#### S2.3 Code implementation and unit tests of CIG-SLM

In the software implementation, _apply_target_mask() constructs a sparse diagonal matrix from the target-cell identity vector and performs receiver-side column gating through W.dot(D). tests/test_masking.py uses a synthetic $4 \times 4$ sparse matrix to verify that columns corresponding to the specified target receptor cells retain their original edge weights, whereas the remaining receptor columns are set to zero, and that right multiplication by $\mathbf{W}\mathbf{D}^{(c)}$ produces a different result from left multiplication by $\mathbf{D}^{(c)}\mathbf{W}$. The tests also cover copying of the unmasked matrix, an invalid mode, and a mismatch in the length of the identity vector.

#### S2.4 Matched LIANA+ bivariate unmasked baseline

The matched LIANA+ bivariate unmasked baseline uses the same expression input, spatial graph, ligand-receptor resource, local statistic, and permutation parameters as the masked mode but does not use $\mathbf{D}^{(c)}$. In this mode, candidate Target_type labels are used only for result alignment and stratification and do not participate in receptor spatial lag calculation. This baseline is an ablation control for receiver-side identity gating in CIG-SLM and is used to assess changes in target-cell communication attribution.

#### S2.5 Local Moran's $R$, local $P$ value, and Sig prop

MaskTalk calculates local bivariate Moran's $R$ for each source cell and obtains a local $P$ value through a local permutation test. Within-sample summary metrics include Mean_Morans_R, Local_sig_cell_count, Total_cell_count, and Local_sig_cell_prop. Local_sig_cell_prop, abbreviated as Sig prop, is defined as the proportion of source cells with a local $P$ value below the prespecified threshold among all tested source cells.

In a grouped multisample design, MaskTalk first aggregates within-sample results at the Subject level and then compares two Conditions. The default direction of the difference is condition2 - condition1. Outputs include Diff_mean_Morans_R, Diff_local_sig_prop, Mann-Whitney U test $P$ values, and Benjamini-Hochberg FDR. MaskTalk first calculates the condition2 - condition1 differences. If min_subject_count is set, records that do not meet the minimum Subject-count requirement in either condition have their difference, $P$ value, and FDR recorded as missing. The Mann-Whitney U test produces a valid $P$ value only when each condition contains at least 2 independent subjects. Because the Visium HD validation contained 1 independent subject per condition, this analysis retained effect-size differences but did not produce valid $P$ values or FDR values.

### S3 Validation analysis design

#### S3.1 Bystander validation

Bystander validation compares the target profiles of the matched unmasked baseline and masked mode across candidate target cell types. In the matched unmasked baseline, candidate target cell types are used only to pair with and stratify the masked outputs and do not enter the spatial adjacency matrix or receptor spatial lag calculation; the curve is therefore expected to lack variation dependent on target-cell identity. In the masked mode, target-cell identity enters $\mathbf{W}^{(c)}$, allowing a target-dependent profile to emerge. Representative axes show typical algorithmic behavior, and the corresponding panels are provided in the Supplementary Figures.

#### S3.2 Mode-specific records

Communication records that pass the prespecified effect-size thresholds are termed threshold-positive LR edges. The mode-specific analysis focuses on masked-specific and unmasked-specific records: masked-specific records reach the prespecified effect-size threshold only in the masked mode, whereas unmasked-specific records reach the threshold only in the matched unmasked baseline. Shared records that reach the threshold in both modes are not included in the main mode-specific analysis but are exported separately; in the all-threshold-positive sensitivity analysis, they are retained in the All masked and All unmasked sets, respectively.

In the Visium HD validation, the summary table first aggregates within-sample results by Subject_ID and then calculates Condition_B - Condition_A. Because the object contains 1 independent subject per condition, Mann-Whitney U test $P$ values and FDR values were not produced. The mode-specific and all-threshold-positive sets in this study were therefore filtered using prespecified effect-size thresholds: |Diff_mean_Morans_R| ≥ 0.05 or |Diff_local_sig_prop| ≥ 0.05, with a valid Mean_Morans_R required in at least one condition. If valid $P$ values are available in future multisubject data, the workflow can use $P$ value and effect-size filters jointly.

The comparison dimensions include target-cell lineage distributions, $|\Delta \text{Moran's } R|$, physical source-target cell distances, and the source-target connection structures of representative ligand-receptor axes. Physical source-target cell distance was calculated from the spatial_um coordinates in AnnData and defined as the Euclidean distance between the spatial centroids of the source and target cell types. This is a record-level descriptive analysis of output differences between the two computational modes.

#### S3.3 All-threshold-positive sensitivity analysis

The all-threshold-positive analysis includes all records that reach the prespecified effect-size thresholds in the two modes and examines whether the trends of the main mode-specific analysis remain directionally consistent in a broader record set. This analysis includes shared records, so the between-mode differences may be weaker than those in the main mode-specific analysis.

#### S3.4 Representative axis network

Representative ligand-receptor axes were selected jointly on the basis of target-cell-dependent profiles, mode-specific effect-size rankings, graphical readability, and consistency with known cell-type and molecular context. These axes were used to show typical algorithmic behavior.

#### S3.5 Public Xenium target-profile range

The public Xenium validation ran matched unmasked and masked inference on a single-sample object and compared Mean Moran's $R$ and Sig prop on spatial contact and kNN graphs. For a fixed combination of Source_type, ligand-receptor pair, and spatial graph type, the target-profile range was defined as the maximum minus the minimum across all candidate target cell types.

Representative Xenium axes were selected from highly ranked target-profile ranges while also considering the consistency of the masked top target with known cell-type and molecular context. The current representative axes include Endothelial MMRN2→CLEC14A, Myoepi_KRT15+ PTN→SDC4, and Endothelial MMRN2→CD93.

### S4 Software reproduction and archiving

#### S4.1 Examples workflow

The examples workflow demonstrates basic MaskTalk usage, including spatial graph construction, communication inference, result summarization, and network- and bubble-plot generation. The workflow comprises 01_build_spatial_graph.py, 02_infer_communication.py, 03_summarize_communication.py, 04_plot_results.py, run_example_pipeline.py, and submit_slurm.sh in the examples directory. This workflow corresponds to Figure 3B-D in the main text.

#### S4.2 demo_validation workflow

The demo_validation workflow reproduces the algorithm validation, including masked/unmasked ablation inference, ablation-result summarization, bystander validation, and mode-specific signal characterization. The workflow comprises 01_ablation_inference.py, 02_ablation_summary.py, 03_bystander_validation.py, 04_signal_characterization.py, run_demo_validation_pipeline.py, and submit_slurm.sh in benchmarks/demo_validation. This workflow corresponds to Figure 2, Figure S1, and Figure S2.

#### S4.3 xenium_janesick workflow

The xenium_janesick workflow reproduces the single-sample public Xenium validation, including AnnData preparation, spatial graph construction, matched unmasked and masked communication inference, target-dependence summarization, and plotting. The workflow comprises 01_prepare_anndata.py, 02_build_spatial_graph.py, 03_infer_communication.py, 04_summarize_validation.py, 05_plot_validation.py, run_xenium_pipeline.py, and submit_slurm.sh in benchmarks/xenium_janesick. This workflow corresponds to Figure S3.

#### S4.4 Tests, API documentation, and Zenodo records

MaskTalk provides lightweight pytest tests, API documentation, a README, example workflows, validation workflows, and Zenodo archives. The API documentation is generated automatically from source-code docstrings and is located at docs/en/api/index.html. Test files include tests/test_graph.py, tests/test_masking.py, and tests/test_statistics.py. The Zenodo data record contains the examples input data, examples processed results, demo_validation input and result data, xenium_janesick processed results, and SHA256 checksum files; the xenium_janesick archive does not contain the raw 10x Genomics Xenium files. Specific DOIs are provided in the Data availability and Code availability sections of the main text.

### Supplementary Figures

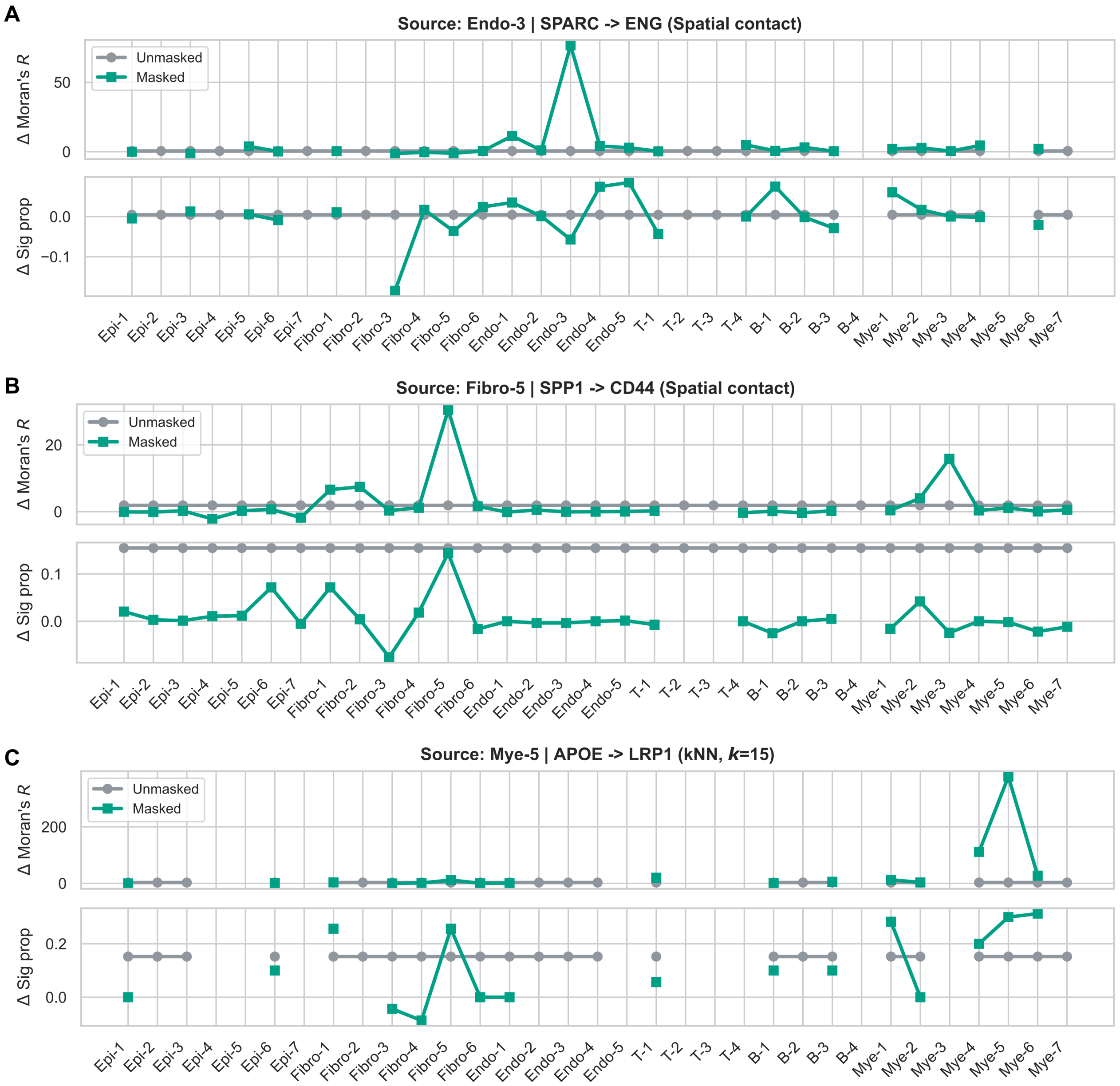


**Figure S1.** Bystander validation results for multiple representative ligand-receptor axes. **(A-C)** Bystander validation results for different representative ligand-receptor axes. The upper panel of each subplot shows $\Delta$Moran's $R$ across candidate target-cell states, and the lower panel shows the corresponding $\Delta$ Sig prop. Gray curves indicate the unmasked mode, and green curves indicate the target-masked mode. The x-axis shows the complete list of cell types in the input object; missing points indicate that no output record was available for the corresponding candidate target cell type. Results are shown for SPARC→ENG with Endo-3 as the source type on a spatial contact graph **(A)**, SPP1→CD44 with Fibro-5 as the source type on a spatial contact graph **(B)**, and APOE→LRP1 with Mye-5 as the source type on a spatial kNN ($k$=15) graph **(C)**.

*Alt text: Bystander validation plots for multiple representative ligand-receptor axes, comparing the target profiles of the unmasked and target-masked modes across candidate target-cell states.*

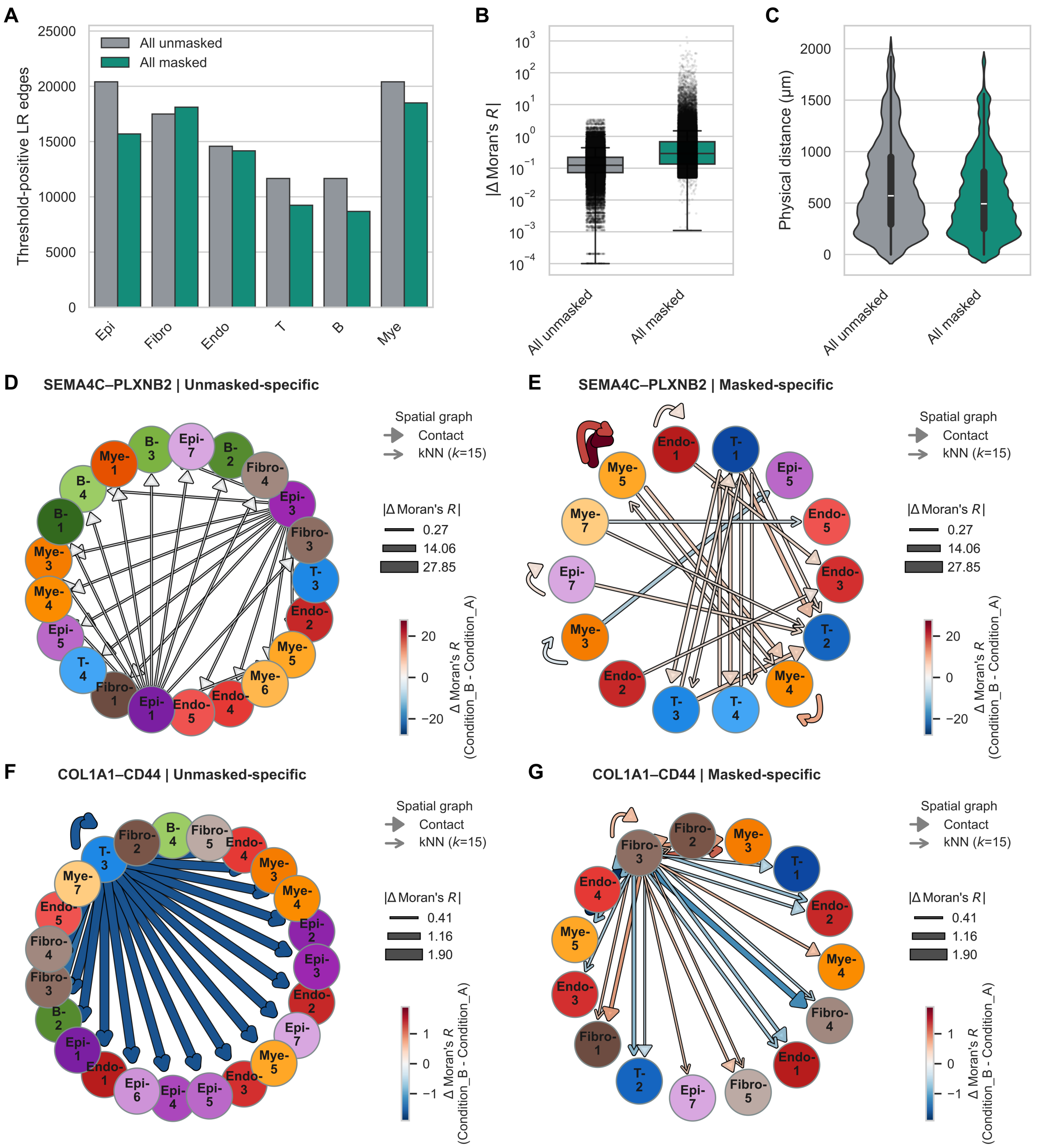


**Figure S2.** Target-masked output characteristics in the complete filtered signal sets and additional representative communication networks. **(A)** Numbers of threshold-positive LR edges among different target-cell lineages for All unmasked and All masked records. All unmasked and All masked denote all communication records that reach the prespecified effect-size thresholds in each mode, including shared signals detected by both modes. Bars summarize the numbers of all filtered communication records in each mode by target-cell lineage. **(B)** Distributions of between-condition effect sizes for All unmasked and All masked records, measured as $|\Delta \text{Moran's } R|$. Boxplots and points show the $|\Delta \text{Moran's } R|$ distributions of all filtered communication records in the two modes. **(C)** Distributions of physical distances between the spatial centroids of the source and target cell types corresponding to All unmasked and All masked records. Violin plots show the physical-distance distributions between the spatial centroids of the source and target cell types corresponding to all filtered communication records in the two modes. **(D-G)** Mode-specific communication networks for additional representative ligand-receptor axes. Nodes denote cell types, and edges denote mode-specific communication records for the corresponding ligand-receptor axis across source-target cell pairs; edge width denotes $|\Delta \text{Moran's } R|$, edge color denotes $\Delta \text{Moran's } R$, and arrow style distinguishes spatial contact and kNN graphs. The unmasked-specific and masked-specific networks for SEMA4C→PLXNB2 are shown in **(D)** and **(E)**, respectively; those for COL1A1→CD44 are shown in **(F)** and **(G)**, respectively.

*Alt text: Sensitivity analysis of the complete filtered signal sets and additional representative networks, comparing the numbers, effect sizes, physical distances, and representative source-target connection structures of All unmasked and All masked signals.*

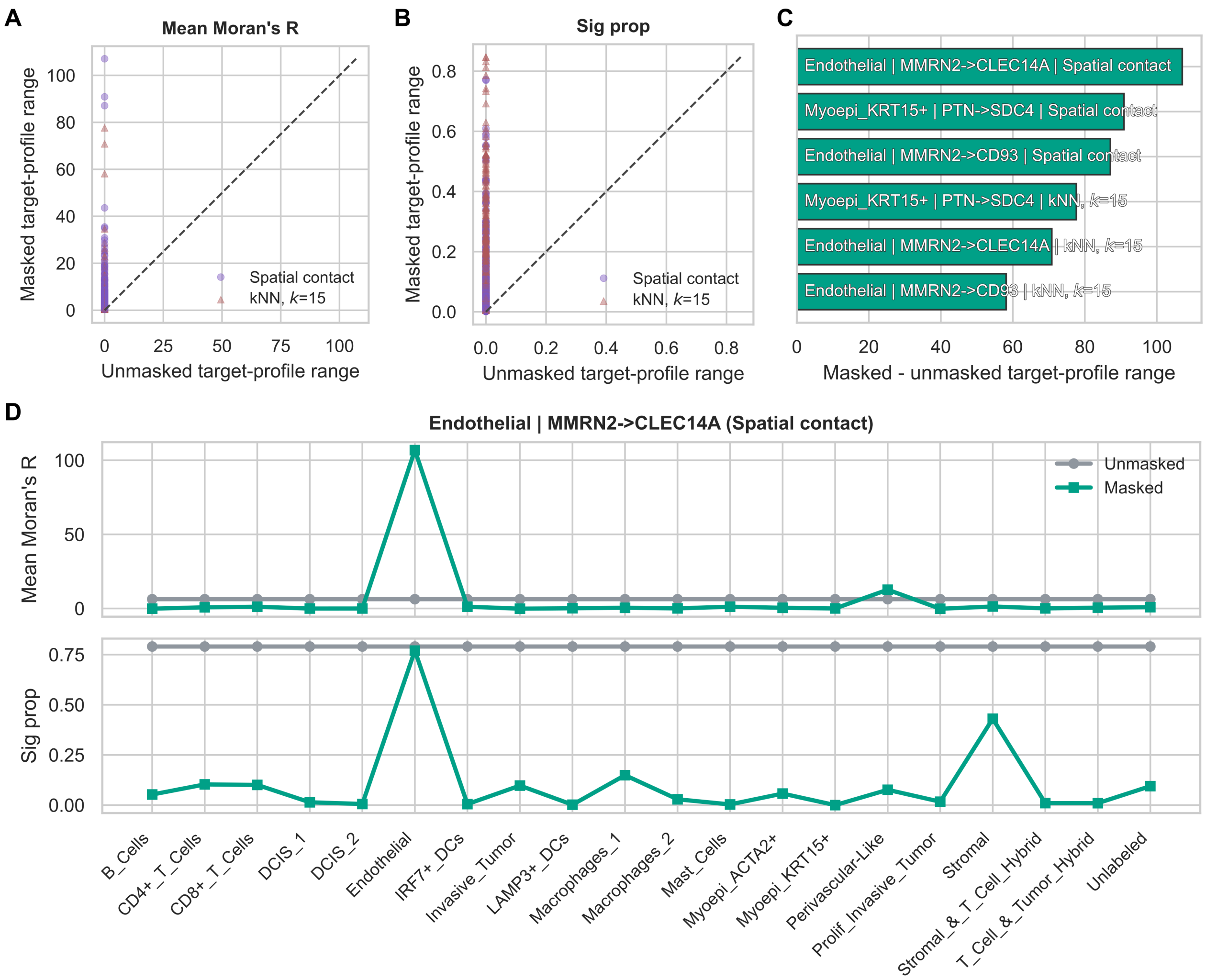


**Figure S3.** Target-masking validation on public 10x Xenium data. **(A,B)** Target-profile ranges from matched unmasked and masked inference in a single public 10x Xenium breast cancer sample. Each point denotes a fixed combination of source cell type, ligand-receptor pair, and spatial graph type. The target-profile range is defined as the maximum minus the minimum across all candidate target cell types for the same combination. The x-axis shows the range from matched unmasked inference, and the y-axis shows the range from masked inference; the dashed line denotes $y = x$. **(A)** Mean Moran's $R$. **(B)** Sig prop. Point color and shape indicate spatial graph type. **(C)** Representative ligand-receptor axes ranked by the masked-minus-unmasked target-profile range. Bar length denotes the difference in the Mean Moran's $R$ target-profile range between masked and matched unmasked inference. Each bar is labeled with the source cell type, ligand-receptor pair, and spatial graph type. **(D)** Target profile of MMRN2→CLEC14A in Endothelial cells on a spatial contact graph. The upper panel shows Mean Moran's $R$ across candidate target cell types, and the lower panel shows the corresponding Sig prop. Gray curves indicate matched unmasked inference, and green curves indicate masked inference. The x-axis shows the complete list of cell types in the Xenium supervised annotation; missing points indicate that no output record was available for the corresponding candidate target cell type.

*Alt text: MaskTalk validation in a single public Xenium sample, showing target-profile range scatterplots, top target-dependent axes, and the target profile of Endothelial MMRN2→CLEC14A.*

### Supplementary Tables

**Table S1. MaskTalk input fields, output fields, and definitions of main statistical metrics**

| Field or metric | Level | Type | Definition | Location |
|---|---|---|---|---|
| adata.X or adata.raw.X | cell × gene | matrix | Expression matrix; the matrix used for LIANA bivariate inference should contain normalized expression, such as log-normalized expression | AnnData |
| adata.obsm[spatial] | cell | matrix | Two-dimensional spatial coordinates of each cell | AnnData |
| Sample_ID | sample | categorical | Tissue sample, section, or field-of-view identifier; graphs for multisample data are constructed separately by this field | adata.obs |

| Field or metric | Level | Type | Definition | Location |
|---|---|---|---|---|
| Subject_ID | independent replicate | categorical | Independent statistical replicate, usually corresponding to a patient in a clinical cohort | adata.obs |
| Condition_ID | group | categorical | Condition or group label used for condition comparison | adata.obs |
| Cell_type | cell | categorical | Cell-type or cell-state annotation used for Source_type and Target_type | adata.obs |
| Graph_key | spatial graph | string | Key of the spatial adjacency matrix in AnnData | adata.obsp / result table |
| Graph_type | spatial graph | string | Readable spatial graph label, such as spatial_contact or spatial_k15 | result table |
| Mask_mode | inference mode | string | masked or unmasked | result table |
| Source_type | cell type | categorical | Source cell type from which ligand expression is obtained | result table |
| Target_type | cell type | categorical | Candidate target cell type used for receiver-side gating or result alignment | result table |
| Ligand_complex | ligand-receptor pair | string | Name of a ligand or ligand complex in the LIANA resource | result table |
| Receptor_complex | ligand-receptor pair | string | Name of a receptor or receptor complex in the LIANA resource | result table |
| Mean_Morans_R | within-sample communication record | numeric | Mean local Moran's *R* between source-cell ligand expression and the receptor spatial lag | communication output |
| Local_sig_cell_count | within-sample communication record | integer | Number of source cells with a local *P* value below the threshold | communication output |
| Total_cell_count | within-sample communication record | integer | Number of tested source cells with a nonmissing local *P* value | communication output |
| Local_sig_cell_prop / Sig prop | within-sample communication record | numeric | Local_sig_cell_count / Total_cell_count | communication output |
| Diff_mean_Morans_R | condition comparison | numeric | Difference in Mean_Morans_R calculated as condition2 - condition1 | aggregate output |
| Diff_local_sig_prop | condition comparison | numeric | Difference in Sig prop calculated as condition2 - condition1 | aggregate output |
| P_value_diff_Morans_R | condition comparison | numeric | Two-sided Mann-Whitney U test *P* value based on subject-level Mean_Morans_R | aggregate output |
| P_adj_FDR_diff_Morans_R | condition comparison | numeric | Benjamini-Hochberg FDR for P_value_diff_Morans_R | aggregate output |
| P_value_diff_sig_prop | condition comparison | numeric | Two-sided Mann-Whitney U test *P* value based on subject-level Sig prop | aggregate output |
| P_adj_FDR_diff_sig_prop | condition comparison | numeric | Benjamini-Hochberg FDR for P_value_diff_sig_prop | aggregate output |
| Diff_communication_score / DeltaComm | visualization summary | numeric | Between-condition communication-difference metric used in network or bubble plots | plot source data |

| Field or metric | Level | Type | Definition | Location |
|---|---|---|---|---|
| Total_perturbation_score / perturbation | visualization summary | numeric | Summary metric used for network edge width or perturbation magnitude | plot source data |

**Table S2. Mapping of main and supplementary figures to figure-source data files**

| Figure | Panel | Content | Figure-source data or source file |
|---|---|---|---|
| Figure 1 | A-E | Schematics of the problem setting, CIG-SLM, local Moran's $R$, and output levels | Author-drawn schematic |
| Figure 2 | A | Representative-axis target profile in demo validation | benchmarks/demo_validation/data/03_bystander_validation/02_bystander_lineplots/ |
| Figure 2 | B-D | Lineage counts, effect sizes, and physical-distance distributions of mode-specific records | benchmarks/demo_validation/data/04_signal_characterization/02_mode_specific_plots/ |
| Figure 2 | E-F | THBS2→NOTCH3 mode-specific networks | benchmarks/demo_validation/data/04_signal_characterization/02_mode_specific_plots/ |
| Figure 3 | A | MaskTalk software workflow schematic | Author-drawn schematic |
| Figure 3 | B-D | Network and bubble plots from the examples workflow | examples/data/04_plot_results/ |
| Figure S1 | A-C | Additional representative axes for bystander validation | benchmarks/demo_validation/data/03_bystander_validation/02_bystander_lineplots/ |
| Figure S2 | A-C | All-threshold-positive sensitivity analysis | benchmarks/demo_validation/data/04_signal_characterization/04_all_significant_plots/ |
| Figure S2 | D-G | Additional mode-specific representative-axis networks | benchmarks/demo_validation/data/04_signal_characterization/02_mode_specific_plots/ |
| Figure S3 | A-D | Target-aware masking validation on public Xenium data | benchmarks/xenium_janesick/data/05_plot_validation/ |

**Table S3. Workflow scripts, outputs, and resource configurations**

| Workflow | Step | Script | Main output | Recommended resources |
|---|---|---|---|---|
| examples | 1 | examples/01_build_spatial_graph.py | examples/data/01_build_spatial_graph/ | 2 cores, 1G |
| examples | 2 | examples/02_infer_communication.py | examples/data/02_infer_communication/ | 8 cores, 17G, array 1-2 |
| examples | 3 | examples/03_summarize_c | examples/data/03_summarize_communication/ | 2 cores, 2G |

| Workflow | Step | Script | Main output | Recommended resources |
|---|---|---|---|---|
| | | ommunication.py | | |
| examples | 4 | examples/04_plot_results.py | examples/data/04_plot_results/ | 2 cores, 1G |
| examples | wrapper | examples/run_example_pipeline.py | examples workflow output | 2 cores, 1G |
| demo_validation | 1 | benchmarks/demo_validation/01_ablation_inference.py | benchmarks/demo_validation/data/01_ablation_inference/ | 8 cores, 18G, array 1-2 |
| demo_validation | 2 | benchmarks/demo_validation/02_ablation_summary.py | benchmarks/demo_validation/data/02_ablation_summary/ | 2 cores, 3G |
| demo_validation | 3 | benchmarks/demo_validation/03_bystander_validation.py | benchmarks/demo_validation/data/03_bystander_validation/ | 2 cores, 1G |
| demo_validation | 4 | benchmarks/demo_validation/04_signal_characterization.py | benchmarks/demo_validation/data/04_signal_characterization/ | 2 cores, 1G |
| demo_validation | wrapper | benchmarks/demo_validation/run_demo_validation_pipeline.py | demo_validation workflow output | 2 cores, 1G |
| xenium_janesick | 1 | benchmarks/xenium_janesick/01_prepare_anndata.py | benchmarks/xenium_janesick/data/01_prepare_anndata/ | 2 cores, 1G |
| xenium_janesick | 2 | benchmarks/xenium_janesick/02_build_spatial_graph.py | benchmarks/xenium_janesick/data/02_build_spatial_graph/ | 8 cores, 2G |
| xenium_janesick | 3 | benchmarks/xenium_janesick/03_infer_communication.py | benchmarks/xenium_janesick/data/03_infer_communication/ | 8 cores, 19G |
| xenium_janesick | 4 | benchmarks/xenium_janesick/04_summarize_validation.py | benchmarks/xenium_janesick/data/04_summarize_validation/ | 2 cores, 1G |

| Workflow | Step | Script | Main output | Recommended resources |
|---|---|---|---|---|
| xenium_janesick | 5 | benchmarks/xenium_janesick/05_plot_validation.py | benchmarks/xenium_janesick/data/05_plot_validation/ | 2 cores, 1G |
| xenium_janesick | wrapper | benchmarks/xenium_janesick/run_xenium_pipeline.py | xenium_janesick workflow output | 2 cores, 1G |

**Table S4. Software environment and tests**

| Item | Requirement or command | Description |
|---|---|---|
| masktalk package | Python >=3.12; masktalk 0.1.1 | Core dependencies are managed by pyproject.toml |
| core dependencies | pyproject.toml | anndata, numpy, pandas, pyarrow, scipy, scanpy, squidpy, liana, networkx, matplotlib, seaborn, statsmodels, and joblib |
| exact reproduction | Python 3.12.13; pip install -r requirements-lock.txt | Install the pinned third-party dependency versions used for the manuscript analyses |
| test extra | pip install -e ".[test]" | Install pytest test dependencies |
| benchmark extra | pip install -e ".[benchmark]" | Install openpyxl for reading the Xenium supervised-annotation xlsx file |
| lightweight tests | pytest -q | Run tests/test_graph.py, tests/test_masking.py, and tests/test_statistics.py |
| API docs | docs/en/api/index.html | Generated automatically from source-code docstrings |

**Table S5. Conceptual positioning of related methods and CIG-SLM**

| Method or framework | Main task | Stage at which cell-type information is used | Spatial or neighborhood modeling | Key difference from CIG-SLM |
|---|---|---|---|---|
| CellChat | Infer cell-population communication networks from ligand-receptor resources | Define source and target cell populations and calculate communication probabilities at the population level | In spatial mode, can integrate spatial distance and filter distant cell-population pairs by the maximum interaction distance of a ligand | Primarily outputs cell-population communication probabilities and networks; CIG-SLM changes the receptor spatial lag in the local statistic |
| CellPhoneDB | Infer cell-population interactions using curated ligand-receptor resources and | Define cell-population pairs, expression means, and candidate- | Microenvironments can constrain the cell-population pairs to be tested | Primarily restricts candidate cell-type pairs; CIG-SLM reconstructs the |

| Method or framework | Main task | Stage at which cell-type information is used | Spatial or neighborhood modeling | Key difference from CIG-SLM |
|---|---|---|---|---|
| | permutation tests | interaction screening | | spatial weight matrix before local-statistic calculation |
| SpatialDM | Identify spatially coexpressed ligand-receptor pairs and local interaction regions using bivariate Moran's statistics | Mainly for result interpretation, cell-type annotation, and visualization | Uses a spatial weight matrix to calculate global and local bivariate Moran's statistics | Local spatial bivariate statistics are the closest component to CIG-SLM; CIG-SLM additionally incorporates target-cell identity into the receptor spatial lag |
| LIANA+ bivariate | Provide ligand-receptor resources and interfaces for multiple spatial bivariate statistics | Can be used for result stratification and input organization for different statistical modules | Uses spatial adjacency and local metrics, including Moran's *R* | MaskTalk reuses its resources and statistical interfaces; CIG-SLM adds receiver-side identity gating and a matched unmasked baseline |
| Squidpy | Provide infrastructure for spatial graphs, spatial autocorrelation, neighborhood enrichment, and ligand-receptor analysis | Used for cluster-level pairs, neighborhood enrichment, or result grouping | Constructs spatial adjacency graphs and provides multiple spatial statistics | Provides the spatial graph and statistical foundation; CIG-SLM addresses receiver-side attribution in directed local ligand-receptor statistics |
| COMMOT | Infer spatial communication flows through collective optimal transport | Cell types can be used for subsequent cluster-level summarization and result interpretation | Constrains optimal-transport flows using spatial distance, communication cost, and molecular competition | Its communication object is an optimal-transport flow; CIG-SLM operates on the receptor spatial lag and local Moran's *R* |

| Method or framework | Main task | Stage at which cell-type information is used | Spatial or neighborhood modeling | Key difference from CIG-SLM |
|---|---|---|---|---|
| SpaTalk | Infer ligand-receptor-target signaling networks by combining spatial neighborhoods and a knowledge graph | Used for sender/receiver cell-type pairs and graph-based cell-pair screening | Quantifies interactions between spatially neighboring cell pairs and links downstream signaling networks | Explicitly distinguishes cell-type pairs but does not define a receiver-side spatial lag gating operator |
| Giotto | Support spatial expression, neighborhood networks, cell interactions, and visualization | Used for enrichment of neighboring ligand-cell-type and receptor-cell-type interactions | Analyzes neighboring cell-type pairs using a spatial proximity network | Performs cell-type-pair neighborhood enrichment; CIG-SLM performs receiver-side gating at the input level of the local statistic |
| stLearn | Analyze trajectories and cell interactions by integrating spatial, morphological, and expression information | Used at the stages of significant hotspots, neighboring pairs, or cell-type-pair summarization | Organizes ligand-receptor hotspots and cell-type pairs using neighboring spots or cells | The target cell type mainly enters hotspot detection, neighbor pairing, or pair-level summarization; CIG-SLM acts before the local statistic |
| MISTy | Explain contributions of spatial context to expression states using multiple views | Can enter as one-hot encodings, cell-type proportions, or stratified-modeling inputs | Represents different spatial scales using intraview, juxtaview, and paraview | A multiview predictive framework; CIG-SLM is an explicitly directed local ligand-receptor statistical operator |
| NICHES | Construct ligand-receptor signal proxies while retaining single-cell and spatial-microenvironment heterogeneity | Used for cell-type grouping, sampling, embedding, and result interpretation | Can construct signal proxies within nearest-neighbor or radius-based neighborhoods | Emphasizes single-cell fidelity and signal proxies; CIG-SLM emphasizes target-cell attribution in the receptor spatial lag |

| Method or framework | Main task | Stage at which cell-type information is used | Spatial or neighborhood modeling | Key difference from CIG-SLM |
|---|---|---|---|---|
| NicheNet | Infer active ligands through a ligand-target prior network | First defines the receiver cell population and then ranks candidate ligands | The original method does not depend on a tissue spatial graph | Focuses on downstream target-gene responses on the receiver side; CIG-SLM focuses on the sources of receptor neighbors in local spatial ligand-receptor statistics |
| Niche-DE | Identify spatial niche-associated differential expression and context-dependent cell-cell interactions | Used for index cell type, niche cell type, and niche-ligand-receptor interpretation | Represents the local environment using effective niches and spatial kernels | A cell-type-specific niche regression method; CIG-SLM is a receiver-side gated receptor spatial lag |
| NCEM | Model the influence of spatial neighborhoods on cellular expression states | Defines index-cell identity and neighborhood context | Represents the local niche using a spatial graph | A spatial-neighborhood expression-prediction framework; CIG-SLM applies receiver-side gating in directed local ligand-receptor statistics |
| SEGCECO | Predict cell-cell communication using graph neural networks and local subgraphs | Mainly used for graph-node interpretation and result evaluation | Focuses on expression-graph or local-subgraph representations rather than explicitly defining a receiver-side receptor spatial lag | A graph-representation-learning prediction method; CIG-SLM does not learn embeddings but explicitly defines a spatial weight operator |
| MESSI | Predict response genes from neighboring-cell signaling features and identify signaling genes | Models are stratified by receiver cell type or state | Uses expression features from neighboring cells as predictive inputs | A supervised predictive model; CIG-SLM is a receiver-side gated extension of local |

| **Method or framework** | **Main task** | **Stage at which cell-type information is used** | **Spatial or neighborhood modeling** | **Key difference from CIG-SLM** |
|---|---|---|---|---|
| | | | | bivariate ligand-receptor statistics |
| MaskTalk / CIG-SLM | Perform target-masked spatial communication inference for a specified target cell type | Incorporates target-cell identity into the receiver-side spatial weight matrix as $\mathbf{D}^{(c)}$ | Defines $\mathbf{W}^{(c)} = \mathbf{W}\mathbf{D}^{(c)}$ before local bivariate Moran's $R$ | Moves target-cell identity into the receptor spatial lag rather than using it only for result grouping or candidate cell-type-pair screening |
